# Metagenomic Detection and Genetic Characterization of Human Sapovirus among Children in Nigeria

**DOI:** 10.1101/2023.12.20.572641

**Authors:** George Etop Uwem, Faleye O. C. Temitope, De Coninck Lander, Agbaje Tunde Sheriff, Ifeorah Maryjoy Ijeoma, Onoja Anyebe Bernard, Oni Igbekele Elijah, Olayinka Adebowale Oluseyi, Ajileye Goodnews Toluwani, Oragwa Obinna Arthur, Akinleye Emmanuel Toluwanimi, Popoola Olubukola Bolutife, Osasona Gideon Oluwadamilola, Olayinka Titilola Olaitan, George Adefunke Oluwadamilola, Muhammad Iluoreh Ahmed, Komolafe Isaac, Adeniji Johnson Adekunle, Matthijnssens Jelle, Adewumi Moses Olubusuyi

## Abstract

Using a metagenomic sequencing approach on stool samples from children with Acute Flaccid Paralysis (AFP), we describe the genetic diversity of Sapoviruses (SaVs) in children in Nigeria. We identified six near-complete genome sequences and two partial genome sequences. Multiple SaV genogroups and genotypes were detected, including GII (GII.4 and GII.8), GIV (GIV.1) and GI (GI.2 and GI.7). Sequence identity and phylogenetic analysis showed that the Nigerian SaVs were related to previously documented gastroenteritis outbreaks associated strains from China and Japan. Minor variations in the functional motifs of the nonstructural proteins NS3 and NS5 were confirmed in the Nigerian strains. To adequately understand the effect of such amino acid changes, a better understanding of the biological function of these proteins is vital. The identification of distinct SaVs reinforces the need for robust surveillance in acute gastroenteritis (AGE) and non-AGE cohorts to better understand SaVs genotype diversity, evolution, and its role in disease burden in Nigeria.

## Introduction

Sapovirus (SaV) infection is a significant public health problem with the virus implicated in acute gastroenteritis (AGE) in humans and animals [1]. The virus has been associated with both outbreaks and isolated cases of AGE among children and adults [2-10]. Sapovirus infections frequently cause diarrhoea and vomiting, which usually last for about a week [11]. However, people exhibiting symptoms for longer than usual and with greater severity have also been documented, particularly in immune-compromised persons [12,13]. Asymptomatic circulation of SaVs has also been reported in children without symptoms of AGE [14,15].

The species *Sapporo virus* belongs to the genus *Sapovirus*, family *Caliciviridae* and was first reported from a diarrheic human stool sample in 1976 using electron microscopy (EM) [16]. The virus is non-enveloped, with a positive-sense, single-stranded RNA genome, which is approximately 7.1 to 7.7 kb in length, containing two open reading frames (ORFs). The large ORF1 encodes a polyprotein which is cleaved by a virus-encoded protease into non-structural proteins (NS1 [p11], NS2 [p28], NS3 [NTpase], NS4 [p32], NS5 [viral genome-linked protein-VPg], and NS6-NS7 [Protease-Polymerase which is further cleaved to form an RNA-dependent RNA-polymerase-RdRp]) and VP1 (major structural protein). The ORF 2 encodes VP2 (minor structural protein) [17-20]. A third ORF has been reported, although its function is currently unknown [1,18].

The most variable (both genetically and antigenically) region of SaVs is the VP1 domain which determines the antigenicity of the virus strains. Classification of SaV is based on complete VP1 amino acid (aa) sequences, with strains having ≥ 57% pairwise VP1 aa identity placed in the same genogroup [21]. Currently, SaVs have been classified into 19 genogroups (GI to GXIX), with four genogroups (GI, GII, GIV and GV) known to infect humans [22-24]. SaVs in other genogroups have been identified in mink (GXII), bats (GXIV, GXVI-GXIX), dogs (GXIII), rodents (GXV), swine (GIII and GV-GXI), and sea lions (GV) [25-27]. The human SaVs are further classified into 17 different genotypes [1], and a recently detected genotype in Peru has been proposed as GII.8 [11,28]. Recombinant SaV strains have been classified as those having a discordant clustering of the VP1 encoding region and the RNA-dependent RNA-polymerase (RdRp) [29], with the RdRp-VP1 junction and the NS3-NS4 junction found to be the two main recombination hotspots [12,30,31].

In Africa, the landscape of circulating human SaV genogroups in recent years has been dominated by GI and GII SaVs [32]. Genogroup V (GV) viruses have been rarely reported in Africa, whereas GIV has been reported in Burkina Faso [33,34] and South Africa [35], where they were identified in up to one-third of infections in patients with gastroenteritis [35,36]. There are currently no published data on SaV infections in Nigeria. In this study, we describe the molecular characterization and genetic diversity of SaV genomes identified in stool samples of children 15 years and below diagnosed with AFP in Nigeria.

## Methodology

### Faecal specimen collection and processing

The faecal samples analysed in this study were collected as part of the National AFP surveillance program in Nigeria. Samples were collected from children aged 15 years and below, diagnosed with AFP in Nigeria in 2020 [37]. These stool samples were collected between January and December 2020 following national ethical guidelines and sent to the WHO National Polio Laboratory in Ibadan, Nigeria.

In this study, 254 archived (−20 freezers stored) poliovirus culture-negative samples from five states in Nigeria (Supplementary Figure 1) were combined into 55 pools by state of collection and the month of sample collection, and subsequently analyzed. Briefly, about 0.5 g of stool was dissolved in 4.5 mL of phosphate-buffered saline (PBS) and 0.5 g of glass beads. After 20 minutes of vortexing, the mixture was subjected to 20 minutes of centrifugation at 3000 rpm. Subsequently, 2 mL of the supernatant was aliquoted into 1 mL cryovials, and stored at -20°C. Thereafter, the stool suspensions were pooled. To make a pool, 200uL faecal suspensions were mixed, with each sample pool containing between 1 to 7 faecal suspensions (Supplementary Table 1). Sample pools were subsequently shipped on ice packs to the University of Leuven, Rega Institute, Laboratory of Clinical and Epidemiological Virology in Belgium.

### Sequencing and Reads processing

The NetoVIR protocol was used to purify virus-like particles (VLPs) from the samples as previously described [38]. Briefly, using a MINILYS homogenizer, faecal suspensions were homogenized for 1 min at 3,000 rpm and filtered through a 0.8 μm PES filter. Free-floating nucleic acids were digested via treatment with a mixture of Benzonase (Millipore, Billerica, MA, USA), (Novagen, Madison, WI, USA), and Micrococcal Nuclease (New England Biolabs, Ipswich, MA, USA). Subsequently, nucleic acid was extracted using the QIAamp Viral RNA Mini Kit (Qiagen, Hilden, Germany) according to the manufacturer’s instructions, but without the addition of carrier RNA. A slightly modified version of the Nextera XT Library Preparation Kit (Illumina, San Diego, CA, USA) was used to prepare the libraries for Illumina sequencing. After that, samples were paired-end sequenced (2x150bp) on an Illumina Novaseq 6000 platform.

Raw reads were processed with the Virome Paired-End Reads (ViPER) pipeline (https://github.com/Matthijnssenslab/ViPER). Using Trimmomatic, the reads were trimmed for quality and adapters [39], and reads mapping to the human genome were removed using Bowtie 2 [40]. Subsequently, the trimmed and filtered reads were *de novo* assembled into contigs using metaSPAdes [41]. The sensitive option in DIAMOND was then used to annotate contigs [42]. KronaTools files were manually inspected to identify all SaV genomes.

### Sapovirus genotyping and Phylogenetic analyses

The SaV sequences generated in this study were aligned with reference human SaV sequences downloaded from GenBank using MAFFT online service [43]. The human calicivirus genotyping tool [44] was used to determine the genogroups and genotypes of each SaV sequence generated in this study. Phylogenetic trees were constructed using MEGA version 11 [45] using the maximum-likelihood method with 1000 bootstrap replications. We aligned each distinct pair of sequences to determine the pairwise identity of the sequences from this study and published reference sequences using the Sequence Demarcation tool [46]. The conserved amino acid motifs for SaV were identified and analyzed using NCBI’s conserved domain database (CDD) [47]. Sequences from this study were also analyzed for recombination events using RDP4 [48].

## Results

Eight (14.5%) of the fifty-five non-polio AFP sample pools tested using deep sequencing contained SaV reads, corresponding to 0.1% to 2.41% of the generated reads (Table 1). Of the eight samples with SaV reads, two sample pools each were from Edo, Abuja, Kaduna and Lagos States (Supplementary Table 1 & Supplementary Figure 1), while no SaV reads were detected in sample pools from Anambra state. We obtained six near-complete genome sequences (SaV-A14-AFP01-NGR-2020, SaV-A16-AFP15-NGR-2020, SaV-A18-AFP18-NGR-2020, SaV-A36-AFP33-NGR-2020, SaV-A143-AFP39-NGR-2020, SaV-A143-AFP46-NGR-2020) with complete coding regions, and two partial genome sequences (SaV-A23-AFP20-NGR-2020 and SaV-A25-AFP10-NGR-2020) with VP1 and VP2 capsid genes but only partial non-structural genes. Complete VP1-based sequence genotyping using the calicivirus typing tool and sequence demarcation tool showed that the eight contigs belonged to three of the four recognized human SaV genogroups (GI, GII and GIV). Further, they belonged to genotypes GI.2 [n=1], GI.7 [n=1], GII.4 [n=3], the newly discovered genotype GII.8 [n=1] and GIV.1 [n=2] (Table 1, Figure 1).

**Table 1:**
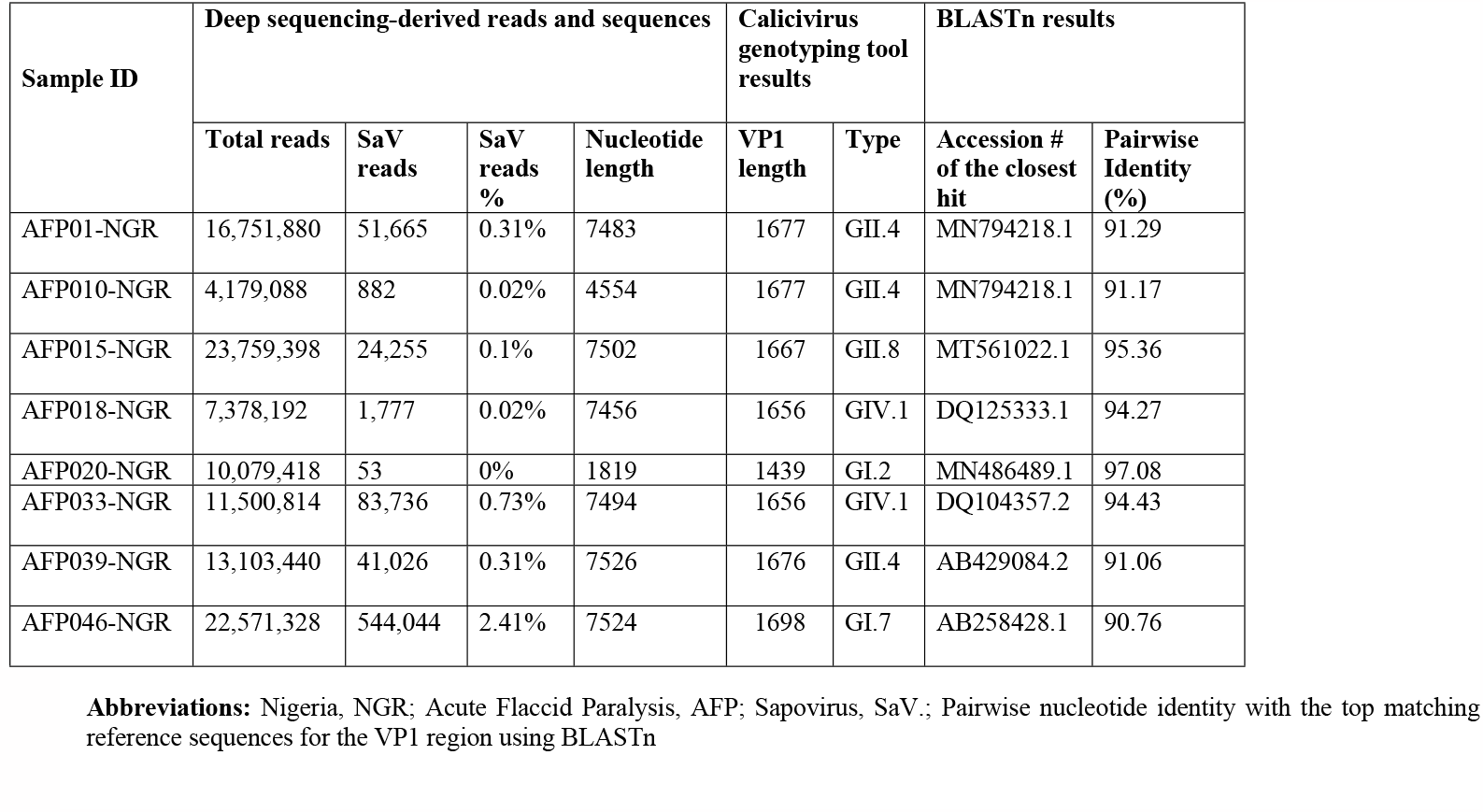
Summary of SaV reads detected in the pooled AFP samples.

**Figure 1:**
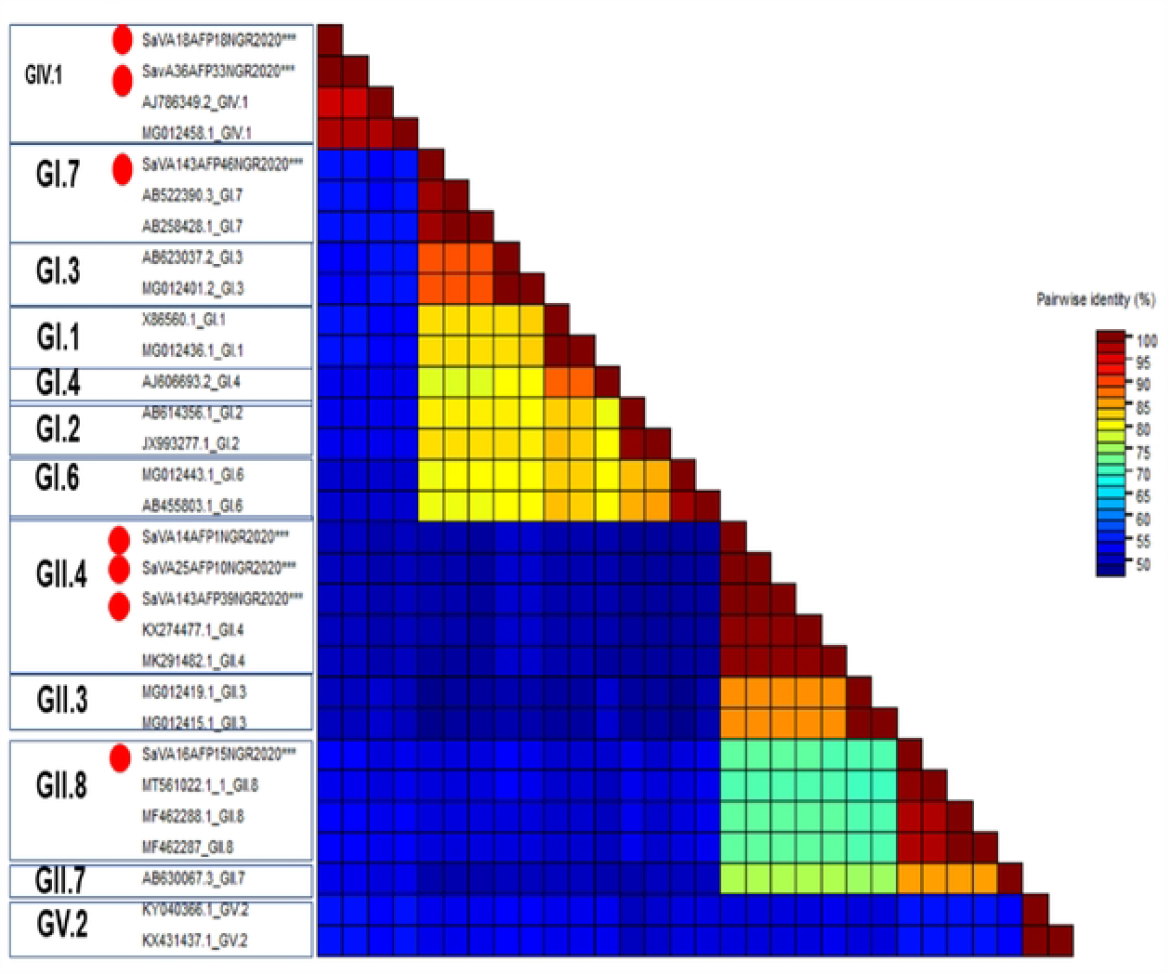
The Calicivirus Sequence Demarcation Tool [46] was used to align the SaV VPI sequences and estimate the pairwise sequence identity between the sequences from this study and existing SaV references. Sequences reported in this study are indicated with a red circle.

Several calicivirus proteins possess conserved motifs and domains responsible for their function. Previously described conserved amino acid motifs in caliciviruses including NS3 [NTpase] (GAPGIGKT), NS5 [Viral genome-linked protein (VPg)] (KGKTK and DDEYDE), protease (GDCG), RNA-dependent RNA-polymerase (WKGL, KDEL, DYSKWDST, GLPSG and YGDD) and VP1(PPG and GWS - have been suggested to be vital in stabilizing P-domain formation in the SaV capsid) [1,18,49]. Analysis of the conserved amino acid motifs of SaV sequences in this study showed minor variations in NS3 and NS5 motifs. In the NS3, all the sequences, irrespective of genogroup, had a GPPGIGKT motif (the A replaced with P), while in the NS5, the KGKTK motif was present in all the sequences except two sequences (SaV-A14-AFP01-NGR-2020and SaV-A143-AFP46-NGR-2020) that had KGKNK motif (Table 2).

**Table 2:**
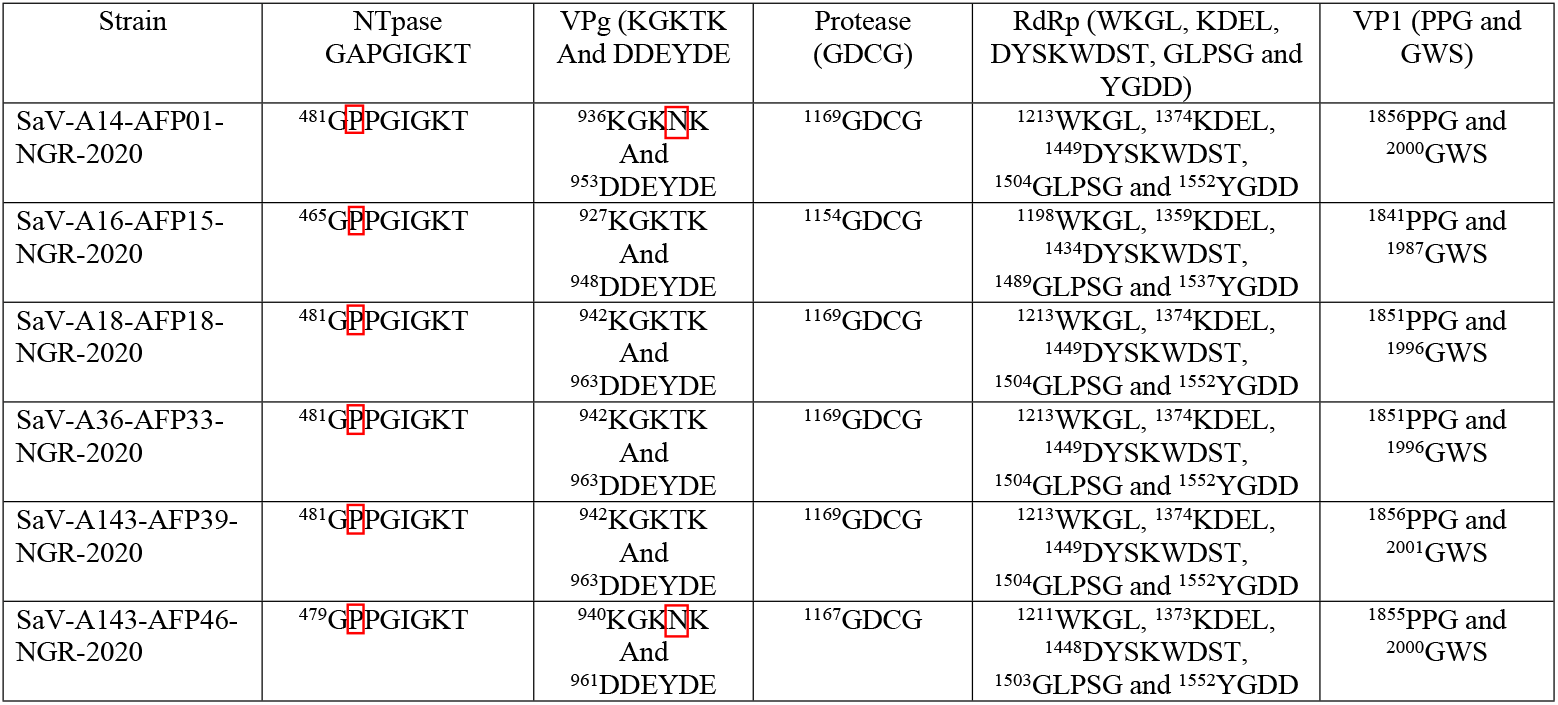
Typical motifs of functional proteins of SaV detected in the pooled AFP samples.

Phylogenetic analysis using the individual genes encoding both the structural (VP1 and VP2) and non-structural proteins (NS1-7) and reference human SaVs showed topological incongruence. Specifically, all the nonstructural genes (Figures 2 and 3) of genomes reported in this study and previously reported reference sequences, including the RdRp gene, clustered into three main genogroups (GI, GII and GV) (Figures 2a-d &3a-d). In contrast, the structural genes (VP1 and VP2) were grouped into four clusters (GI, GII, GIV and GV) (Figures 4b-c). All GIV nonstructural genes were consistently found among GII clusters, while their structural genes were in a group independent of GII. Interestingly, the GIV sequences in this study clustered independently from previously documented strains from Asia and North America. The GII.8 detected in this study clustered with a novel variant of the GII.8 genotype, which was associated with an outbreak of SaV among primary school students in Shenzhen city, China, in 2019 [50]. However, human SaV sequences detected in this study did not contain any significant recombination breakpoints, according to RDP4 sequence analysis.

**Figure 2.**
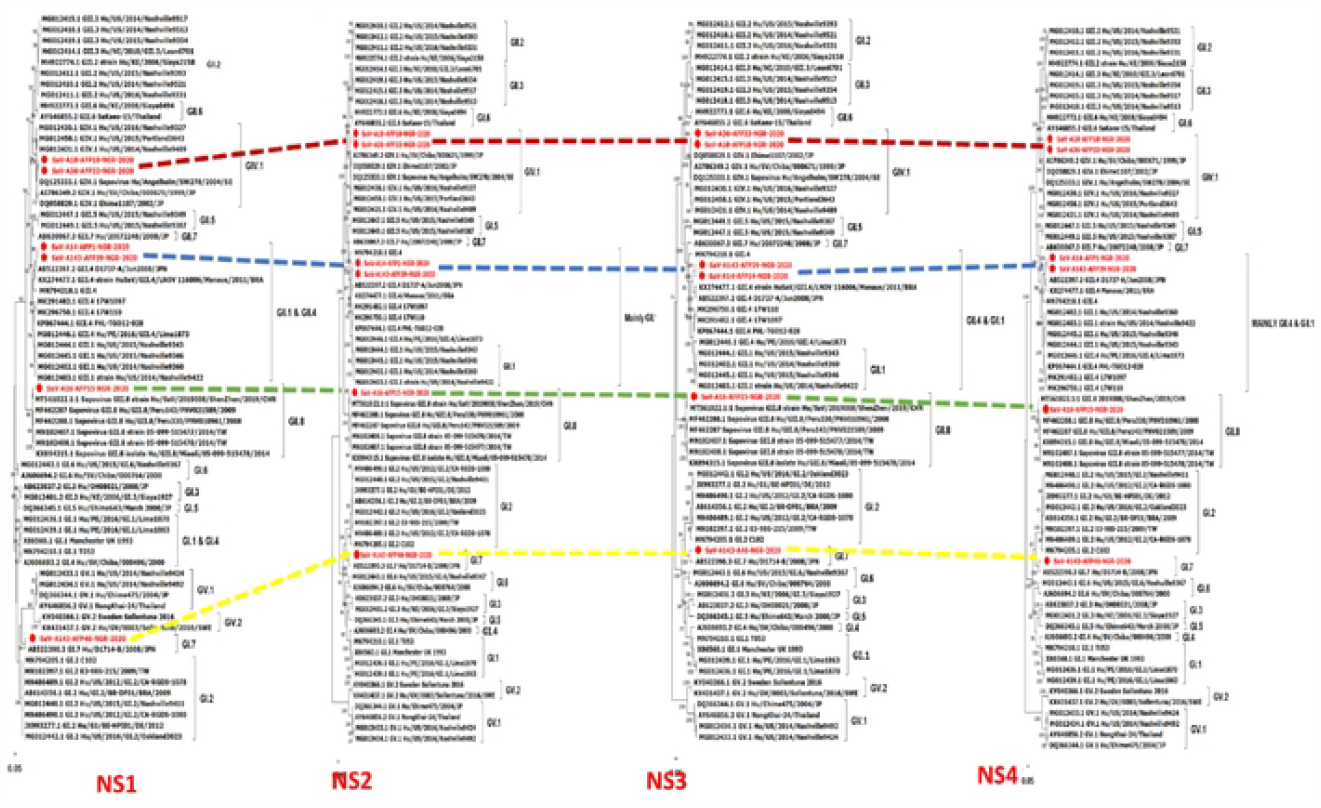
Maximum likelihood trees ofthe NS1-4 genes. The trees were constructed based on the full-length amino acid sequences of (a) the p11 (NS I) protein, (b) the p28 (NS2) protein, (c) the NTpase (NS3), and (d) the p32 (NS4). Bootstrap support values greater than 50 are shown. Sequences report in this study are highlighted in red.

**Figure 3.**
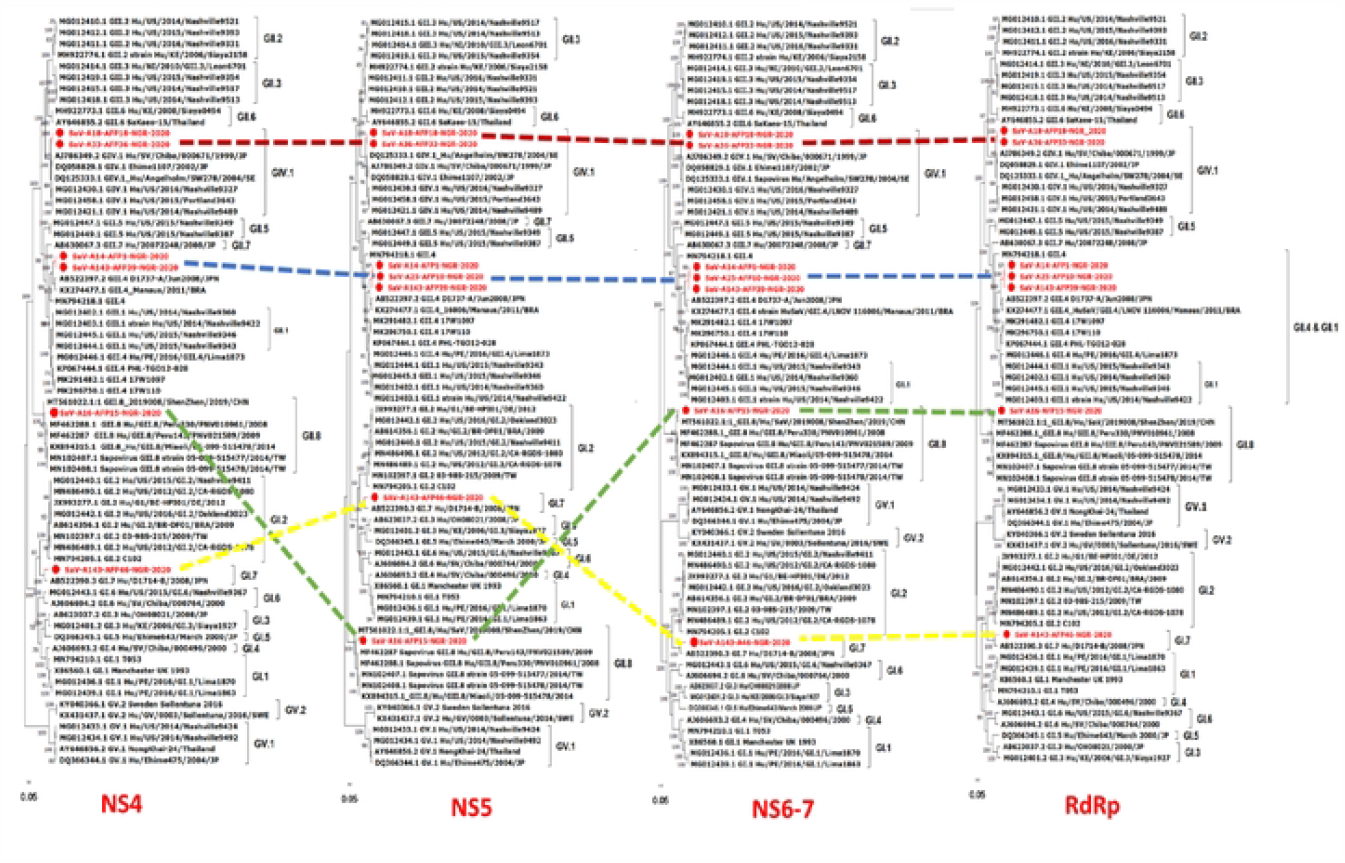
Maximum likelihood trees of NS4-7 and RdRp gene. The trees were constructed based on the full-length amino acid sequences of (a) the p32 (NS4) protein, (b) the viral genome-linked protrein (NS5), (c) the Protease Polymerase (NS6-7), and (d) the RdRp. Bootstrap support values greater than 50 are shown. Sequences report in this study are highlighted in red.

**Figure 4.**
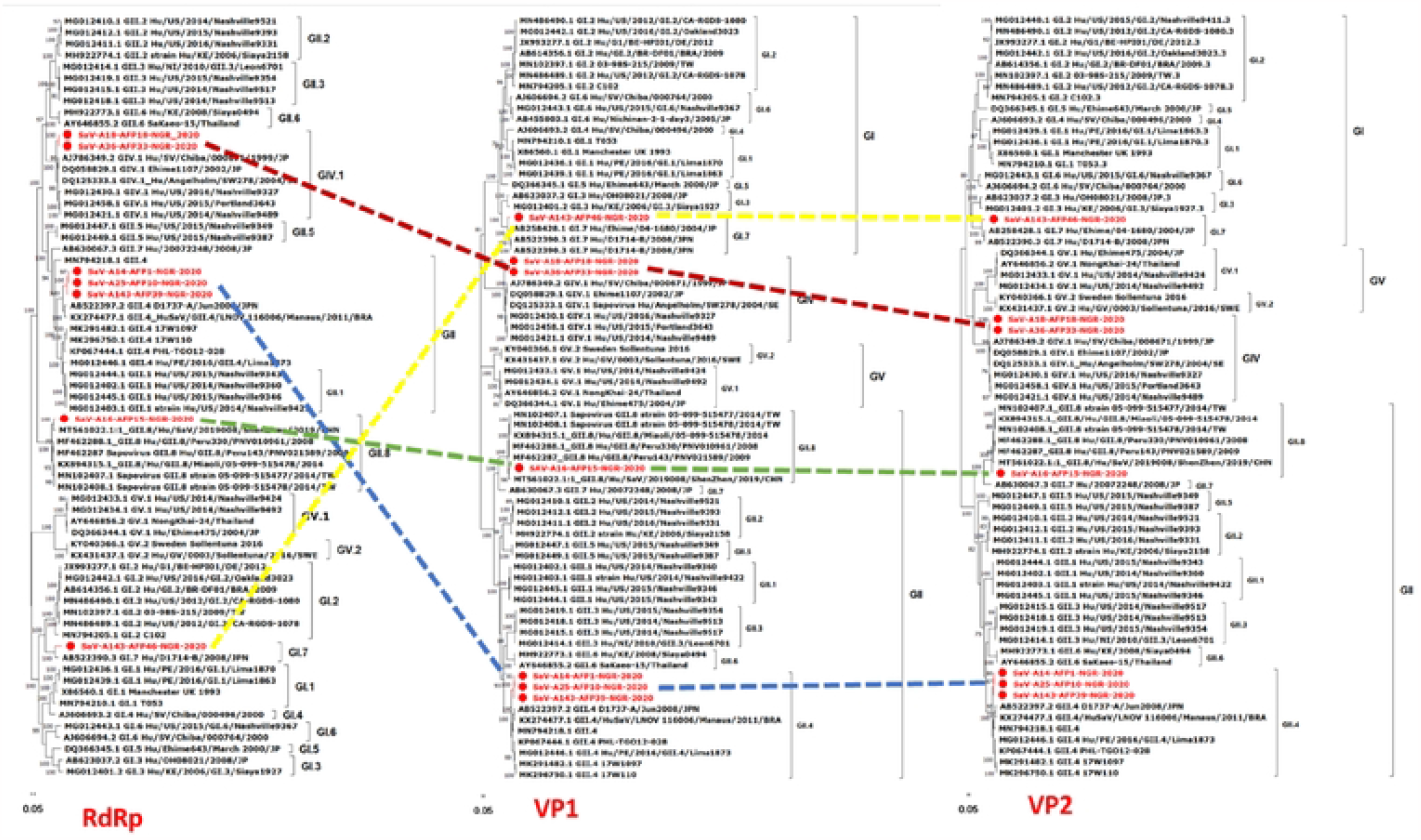
Maximum-likelihood trees of SaV genes. The trees were constructed based on the full-length amino acid sequences of (a) the RdRp, (b) the capsid protein (VP1) and (c) the small basic protein (VP2). Bootstrap support values greater than 50 are shown. Sequences report in this study are highlighted in red

## Discussion

Without a doubt, the global health community has made significant investments and taken targeted actions to address the primary causes of child death through high-impact interventions such as access to nutrition, safe water, sanitation, and vaccination. Malnutrition and diarrheal diseases, on the other hand, continue to be among the top causes of death among children [51,52]. In Nigeria, there is a dearth of information on SaV genetic diversity, epidemiology and evolution [53]. In the present study, we describe six near-complete genome sequences (all having complete coding regions) and two partial genome sequences from children with AFP. This is the first detection of human SaVs in Nigeria. Interestingly, multiple genotypes were detected, indicating the circulation of various strains in Nigeria. Specifically, we document the presence of genogroups GII (GII.4 and GII.8), GIV (GIV.1) and GI (GI.2 and GI.7) in Nigeria.

All the identified human SaVs, irrespective of genotype, had amino acid substitution A482P in the NS3 motifs (Table 2). A similar motif has been reported in SaVs from pigs [25]. Since many caliciviruses, including SaVs, are difficult to grow in cell culture, studying the biological function of their nonstructural proteins remains challenging. However, few studies have elucidated the role and activities of the polymerase and protease (3C-like protease (NS6) and the 3CD-like protease-polymerase (NS6-7) [18,54-56]. Interestingly, mutational analysis of the RdRp conserved GDD amino acid motif from a calicivirus-rabbit hemorrhagic disease virus (RHDV), showed that substitution of the RHDV 3D^pol^ 1605 aspartate residue by asparagine, glycine or glutamate residues resulted in a complete loss of enzymatic activity [57]. Understanding the biological functions of various SaV proteins and the role of various amino acid substitutions in the evolution of the virus is needed to understand the potential implications of newly observed mutations.

Phylogenetically, the Nigerian SaVs were related to previously reported SaV reference strains. While GIV sequences in this study formed small sub-clusters independent from previously documented strains from Asia and North America, the GII.8 in the study was 95.4% similar and clustered with a novel variant of the GII.8 genotype, which was associated with an outbreak of SaV among primary school students in Shenzhen city, China, in 2019 [50]. The position and length of the ORFs, VP1 and VP2, of the Nigerian GII.8 strains were identical to those of the Shenzhen strain. Remarkably, the GI.7 strain from this study was more than 90% similar to GI.7 strains from Japan that were associated with gastroenteritis outbreaks linked to the consumption of contaminated shellfish [58]. Robust surveillance of SaV among AGE and non-AGE cohorts in Nigeria is needed to better understand the genotype diversity, evolution and probable disease association of this virus in the country. Of note, the RDP4 findings and the phylogenetic tree structure did not provide adequate support to classify any of the Nigerian SaVs as recombinant strains.

In conclusion, we describe six near-complete and two partial SaV genome sequences. This is the first report of human SaVs in Nigeria. Hence, the data described here could serve as references to help develop tools to enhance surveillance of and improve epidemiological information about SaVs in Nigeria and Africa at large, where only short genome regions have been reported. Further, understanding the evolutionary dynamics of SaV, especially the nonstructural proteins would be vital to fully delineate the role of amino acid substitutions in SaV evolution and genetic diversity. This would make their nonstructural proteins desirable targets for developing therapeutics to treat human calicivirus infections.

## Data Availability Statement

The Sapovirus nucleotide sequences identified in this study have been deposited in GenBank under the accession number OR837774 - OR837781. Raw sequence data have also been uploaded under the Bioproject number PRJNA1043841.

## Author’s contributions

Sample selection and processing - GEU, ATS, OAO, OIE, AET, POB, OOT, and GAO; Study design – GEU, FOCT, AAJ, MJ and AMO; Laboratory analysis – DCL and MJ; Data analysis – DCL, and GEU; Writing - initial draft of the manuscript – GEU; Writing – Review and editing – FOCT, DCL, ATS, OAB, OAO, AGT, OOA, OGD, AIM, KIO, AAJ, MJ and AMO; Supervision – AAJ, MJ and AMO. All the authors have read and approved the final version of the manuscript before submission.

